# Molecular mechanisms of Evening Complex activity in Arabidopsis

**DOI:** 10.1101/584854

**Authors:** Catarina S. Silva, Aditya Nayak, Xuelei Lai, Veronique Hugouvieux, Jae-Hoon Jung, Agnès Jourdain, Irene López-Vidriero, Jose Manuel Franco-Zorrilla, François Parcy, Kishore Panigrahi, Philip A. Wigge, Max Nanao, Chloe Zubieta

## Abstract

The Evening Complex (EC), composed of the DNA-binding protein LUX ARRHYTHMO (LUX) and two additional proteins, EARLY FLOWERING 3 (ELF3) and ELF4, is a transcriptional repressor complex and a core component of the plant circadian clock. In addition to maintaining oscillations in clock gene expression, the EC also participates in temperature and light entrainment and regulates important clock output genes such as *PHYTOCHROME INTERACTING FACTOR 4* (*PIF4*), a key transcription factor involved in temperature dependent plant growth. These properties make the EC an attractive target for altering plant development through targeted mutations to the complex. However, the molecular basis for EC function was not known. Here we show that binding of the EC requires all three proteins and that ELF3 decreases the ability of LUX to bind DNA whereas the presence of ELF4 restores interaction with DNA. To be able to manipulate this complex, we solved the structure of the DNA-binding domain of LUX bound to DNA. Using structure-based design, a LUX variant was constructed that showed decreased *in vitro* binding affinity but retained specificity for its cognate sequences. This designed LUX allele modulates hypocotyl elongation and flowering. These results demonstrate that modifying the DNA-binding affinity of LUX can be used to titrate the repressive activity of the entire EC, tuning growth and development in a predictable manner.

**Significance Statement:** Circadian gene expression oscillates over a 24 hr. period and regulates many genes critical for growth and development. In plants, the Evening Complex (EC), a three-protein repressive complex made up of LUX ARRYTHMO, EARLY FLOWERING 3 and EARLY FLOWERING 4, acts as a key component of the circadian clock and is a regulator of thermomorphogenic growth. However, the molecular mechanisms of complex formation and DNA-binding have not been identified. Here we determine the roles of each protein in the complex and present the structure of the LUX DNA-binding domain in complex with DNA. Based on these data, we used structure-based protein engineering to produce a version of the EC with altered *in vitro* and *in vivo* activity. These results demonstrate that the EC can be modified to alter plant growth and development at different temperatures in a predictable manner.

## Introduction

The circadian clock provides endogenous rhythms that allow plants to anticipate and react to daily environmental changes. Many processes such as photosynthesis and growth occur in a rhythmic manner over a 24-hour cycle (1–3). These circadian rhythms persist even in the absence of light/dark cues due to internal repeating oscillations of core clock genes that in turn modulate gene expression patterns of many different output pathways (4). In Arabidopsis, the circadian clock consists of three main interacting transcription-translation feedback loops: the morning, central and evening loops. Components of these interlocking feedback loops repress each other’s expression resulting in rhythmic gene expression over a 24-hr period (for review see,(3, 5–7)). The Evening Complex, composed of LUX, ELF3 and ELF4, is a core component of the circadian clock (8–12). The expression patterns of the three genes overlap, peaking at dusk. Thus, the EC has maximum activity at the end of the day and early night, acting to repress expression of the circadian morning loop genes *PSEUDORESPONSE REGULATOR 7* and *9 (PRR7* and *9)*, the central loop gene *CCA1* and the evening loop genes *GIGANTEA (GI)* and *LUX* itself (12–15).

Loss-of-function mutations in *elf3, elf4* or *lux* give rise to arrhythmic circadian outputs with alterations in many developmental pathways (9, 16–18). This results in phenotypes including elongated hypocotyls and early flowering regardless of day length or ambient temperature (9, 12, 16–19). In addition, natural variation in EC components *ELF3* and *LUX* has been shown to give rise to altered thermal responsive growth not only in Arabidopsis but also in crop plants (20–23). Lack of thermoresponsiveness, elongated hypocotyls and early flowering in EC mutants is hypothesized to be due in large part to misregulation of the circadian output pathway involving the bHLH transcription factor *PHYTOCHROME INTERACTING FACTOR 4* (*PIF4)* (15), a master regulator of cell elongation, thermoresponsive growth and the shade avoidance response (19, 24–28).

The repressive regulatory activity of the EC is temperature dependent, making it a node that integrates both circadian gene regulation and environmental information to control growth and developmental pathways in the plant (14, 20, 29). Extensive ChIP-seq experiments performed at different temperatures demonstrated that the binding sites for LUX, ELF4 and ELF3 extensively overlap and that the strength of binding for the EC is dependent on temperature, with weaker binding of the complex at higher temperatures (15). This raises the intriguing possibility that the EC can be used to alter plant growth and development and may be an attractive target for crop improvement (29). The underlying mechanisms that determine EC complex formation and DNA-binding, however, remained to be elucidated. Here, we determine the role of each protein in EC formation, address the molecular determinants of DNA-binding affinity and specificity and demonstrate that alterations in the DNA-binding affinity of LUX can predictably alter EC function *in planta*.

## Results

### Role of LUX, ELF3 and ELF4 in complex formation and DNA-binding

Of the three EC proteins, only LUX is able to directly bind DNA, however complex formation is necessary for full EC activity based on the similar phenotypes of *elf3*, *elf4* and *lux* mutants (9, 16–18). In order to investigate the roles of the different EC components in protein-protein interactions and DNA-binding, *in vitro* refolding of the full EC and subcomplexes was performed due to low solubility of ELF3 under native conditions. Step-wise dialysis against decreasing urea concentrations was used to form the respective complexes. To confirm production of active EC and to assess the DNA-binding affinity of the complex, EMSAs were performed using a 36 base pair (bp) fragment from the *PRR9* promoter containing a previously characterised LUX binding site (LBS) (10). As shown in Figure 1A, ELF3 and ELF4 alone did not interact with DNA, as expected, since neither protein is predicted to have a DNA-binding domain. Addition of ELF4 to a solution containing LUX had no effect on LUX binding. However, titrating in ELF3 while keeping LUX and ELF4 concentrations constant resulted in the disappearance of the LUX-DNA band and the appearance of a higher molecular weight band corresponding to the EC bound to DNA (Figure 1A, *left*). Interestingly, without ELF4 present, LUX-ELF3 exhibited highly attenuated DNA-binding, with the appearance of a free DNA band that increases in intensity with increasing ELF3 concentration. No higher molecular weight bands were observed with LUX-ELF3 alone (Figure 1A, *right*). These results suggest that ELF3 modulates efficient binding of LUX to DNA and that ELF4 is able to restore DNA-binding competency to the complex (Figure 1B).

**Figure 1.**
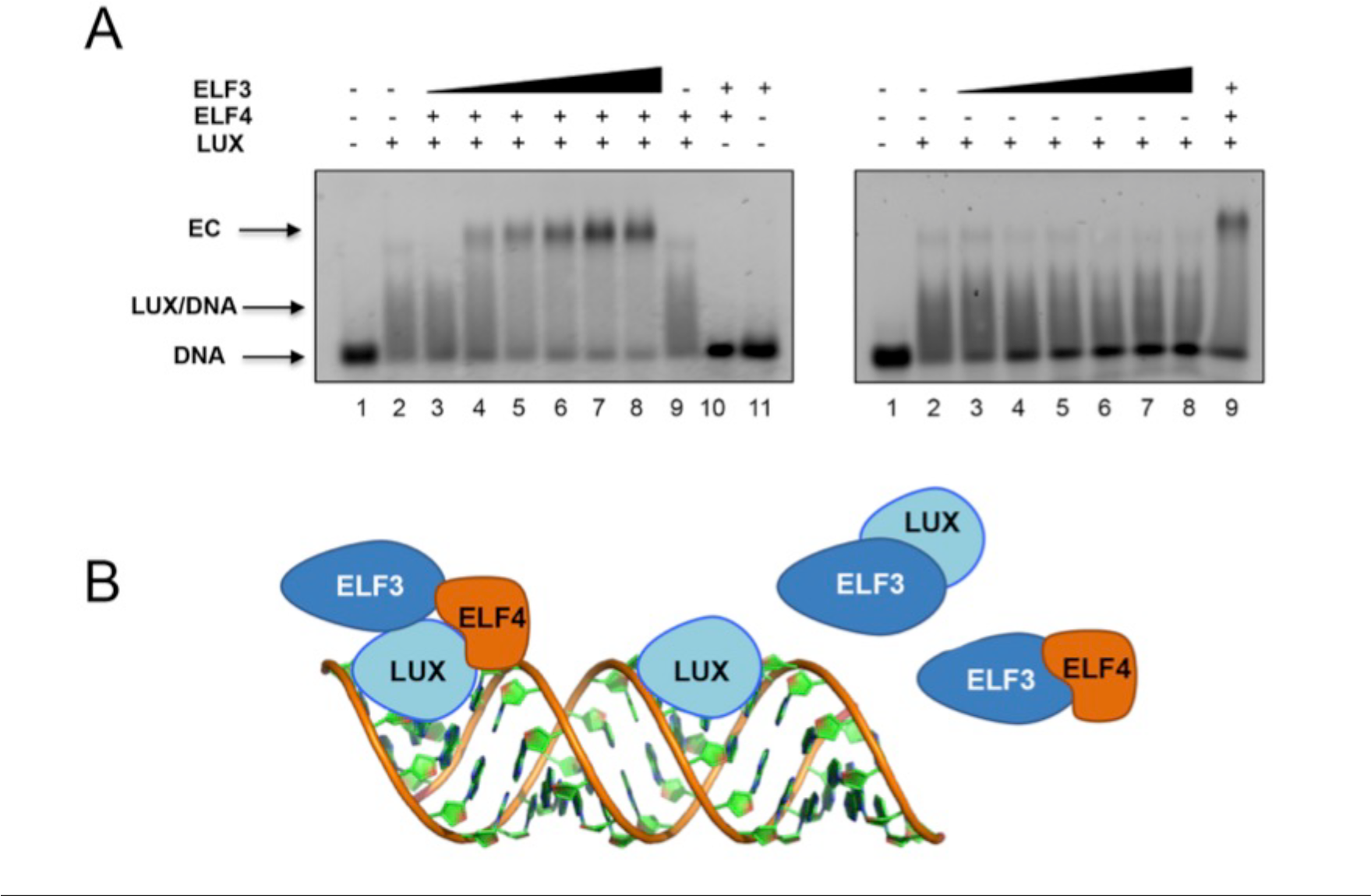
EC and subcomplexes interactions with DNA. **A**. EMSAs of LUX-ELF3 and the EC, in 2% agarose gels. DNA concentration was 30 nM. (Left) Reconstitution of the EC with LUX and ELF4 concentrations held constant at 200 nM and 1000 nM, respectively and increasing ELF3 concentrations (220 nM, 450 nM, 890 nM, 1.3 µM, 1.8 µM and 2.2 µM). (Right), LUX-ELF3 interactions, with LUX concentration kept at 200 nM. With increasing ELF3 concentration, the free DNA band increases in intensity. **B.** Schematic depiction of LUX, ELF3 and ELF4 interactions and DNA-binding.

### LUX binds DNA with high affinity independently of the EC

In order to determine whether LUX alone was sufficient to confer DNA-binding affinity and specificity, we analysed the DNA-binding activity of LUX using protein-binding microarrays (PBM) and the full-length protein (LUX^FL^) and DNA-binding MYB domain (residues 139-200, LUX^MYB^) tagged with an N-terminal maltose binding protein. Experiments were performed and analysed as previously described (30, 31). LUX^FL^ yielded over 100 high affinity binding 8-mers with E-scores over 0.45, indicative of tight binding. Most motifs correspond to variations of the sequence “AGAT(A/T)CG” as previously determined *in vivo*(10) (Figure 2A). The isolated DBD LUX^MYB^ bound with lower affinity producing consensus motifs with E-scores mainly below 0.40, with only two 8-mers identified with E-scores above 0.45. (SI Figure 1).

**Figure 2.**
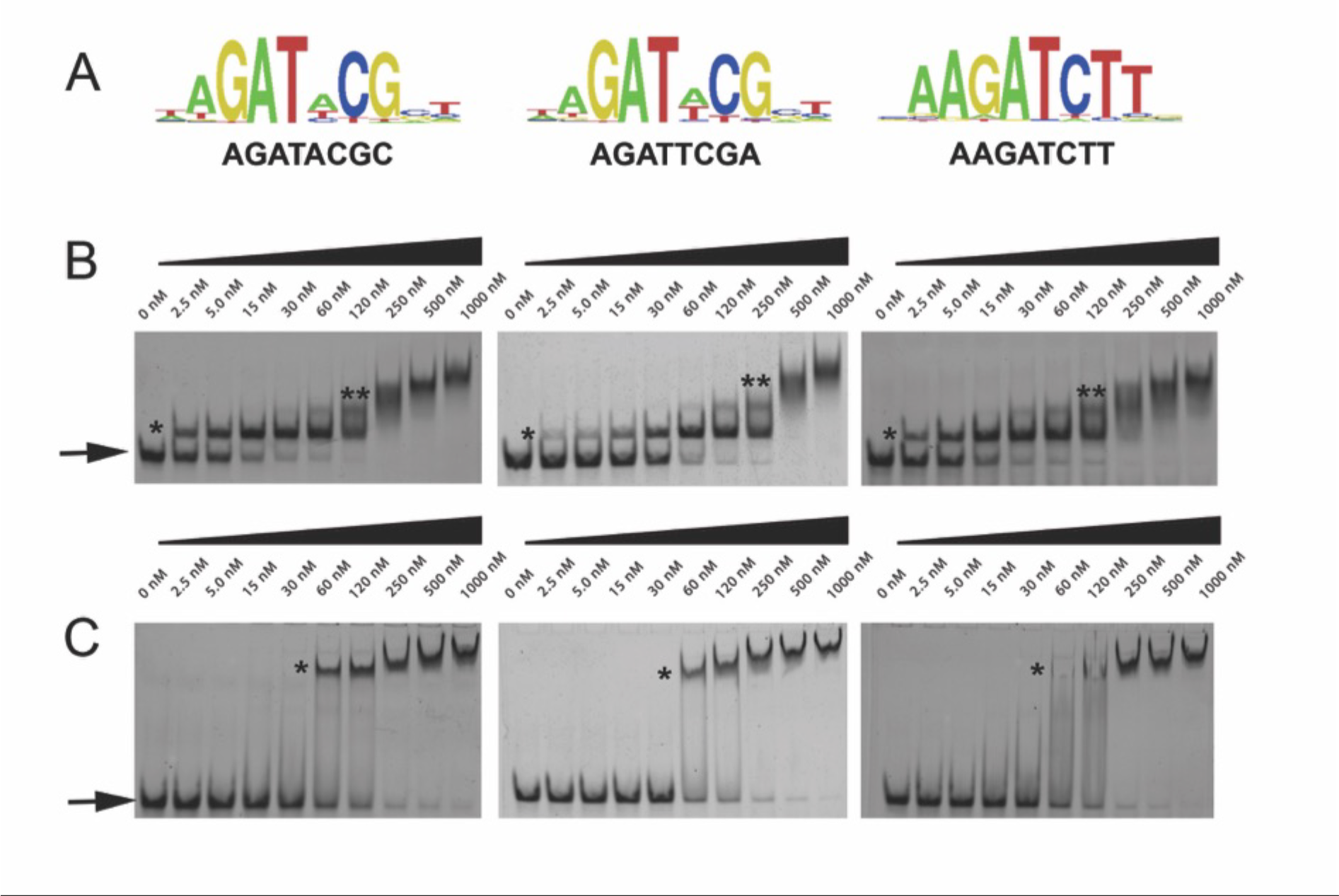
LUX-DNA interactions. **A.** High scoring protein binding microarray-derived logos for LUX. Three logos are presented including the LBS consensus (left), the *PRR9* promoter LBS sequence (middle) and a high scoring PBM sequence (right). **B**. Representative gel electrophoretic mobility shift assays (EMSAs) for LUX^MYB^. DNA concentration was constant with protein concentration increasing from 0 to 1000 nM. The DNA sequences used correspond to the above motifs in (**A**). Free DNA is indicated by an arrow and protein-DNA complexes are indicated with stars. One star corresponds to one molecule of protein bound, two stars indicates multiple non-specifically bound protein molecules at high protein concentrations. **C.** Representative EMSA for LUX^FL^, labelled as per (**B**).

As LUX has only a single MYB domain and neither ELF3 nor ELF4 directly binds DNA, the absolute binding affinities of untagged LUX^FL^ and LUX^MYB^ were assayed to determine whether this single domain would be able to target the EC to its cognate binding sites. To confirm the affinity of LUX-DNA interactions, DNA sequences with variations of the LBS were tested using electrophoretic mobility shift assays (EMSAs) with varying protein concentrations. Surprisingly, LUX^MYB^ exhibited higher affinity compared to the full-length protein for all DNA probes tested with *K*_*d*_’s ranging from 6.5 nM to 43 nM (Figure 2B and Table 1). The full-length protein exhibited lower affinity over the sequences tested, with *K*_*d*_’s in the 90-180 nM range (Figure 2C and Table 1). All *K*_*d*_ measurements were performed on untagged proteins, unlike the PBM experiments. The PBM result indicating a lower affinity of LUX^MYB^ for DNA is thus likely due to the N-terminal MBP fusion, which may occlude the DNA-binding site and suggests that large protein fusions close to the N-terminus of the MYB domain negatively impacts DNA-binding. These data demonstrate that LUX is able to bind with high affinity to its cognate sites in the low nanomolar range and that this tight binding is likely sufficient to target the entire EC to its cognate sites.

**Table 1.**
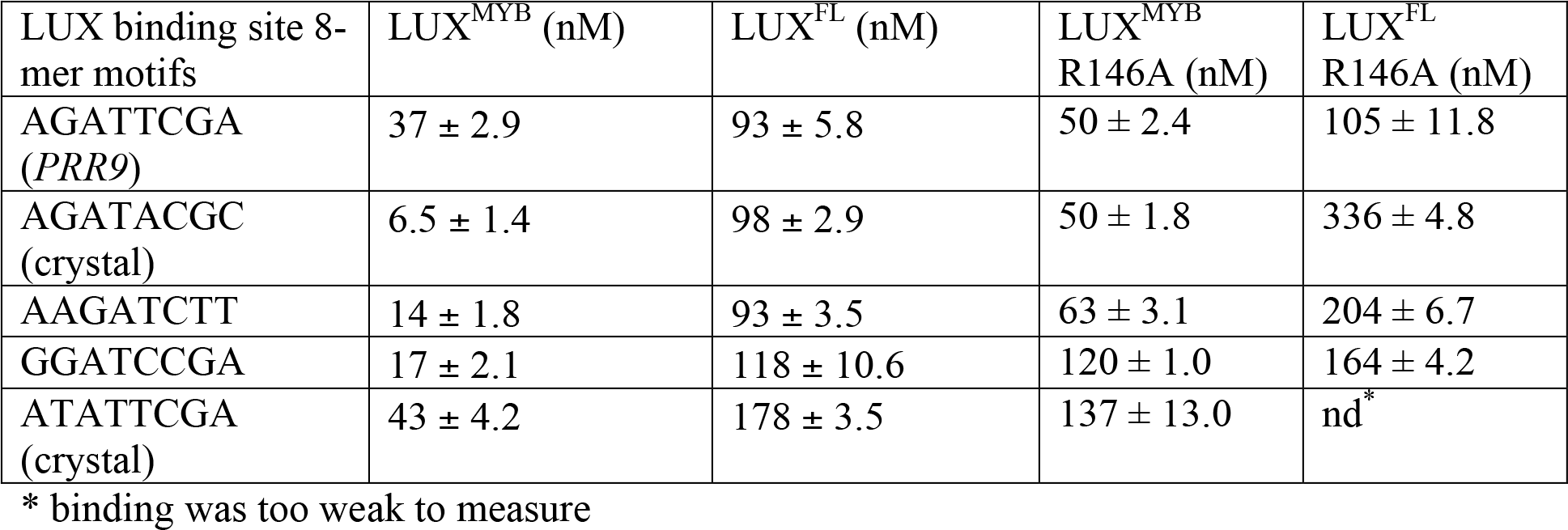
DNA-binding affinities of LUX^MYB^ and LUX^FL^ and respective mutants.

### The GARP family signature motif in LUX is required for base read-out

Having determined the *in vitro* binding specificity and affinity of LUX and LUX^MYB^ with DNA, we sought to reveal the molecular determinants for DNA-binding. LUX^MYB^ was crystallized and the atomic structure determined in complex with a 10-mer dsDNA, 5’-TA<u>GATACG</u>CA containing the core binding motif (underlined) determined from the PBM experiments and a second DNA sequence 5’-TA<u>TATTCG</u>AA which lacks the highly conserved guanine at the beginning of the LBS and replaces the adenine with a thymine at position 4, a conservative change in the LBS consensus sequence GAT(A/T)CG (SI Table 1). For both structures, LUX^MYB^ adopts a classic three-helix bundle conformation characteristic of MYB domains (Figure 3). The MYB hydrophobic core usually consists of three regularly spaced residues, most often tryptophans, with a spacing of 18 or 19 amino acids (32). In LUX^MYB^, however, the second and third tryptophan residues are replaced by a proline (Pro171) and leucine (Leu192) based on structural alignments. Proline at position 171 creates a tight turn before helix 2 and brings the helix in close proximity to the DNA. A proline at this position is also conserved in other plant MYB proteins such as the structural characterized transcription factor, AtARR10 (Figure 3B). Interestingly, in the *lux-4* null mutant, Pro171 is replaced by a leucine residue, suggesting the tight turn before helix 2 is required for proper interaction of the protein with DNA. The hydrophobic core in LUX^MYB^ is stabilized by additional hydrophobic interactions including edge-to-face interactions of Phe157 (helix 1) and Tyr195 (helix 3), π stacking of Trp149 (helix 1) with His191 (helix 3) and edge-to-face interactions of His191 and Phe157 (Figure 3A). The protein sequesters DNA primarily through helix 3 that lies in the major groove and contains a plant-specific GARP family signature motif, SH(A/L)QK(F/<u>Y</u>) (16). Examination of the electrostatic surface of the protein demonstrates a highly electropositive face that acts as the main DNA-binding surface (Figure 3A).

**Figure 3.**
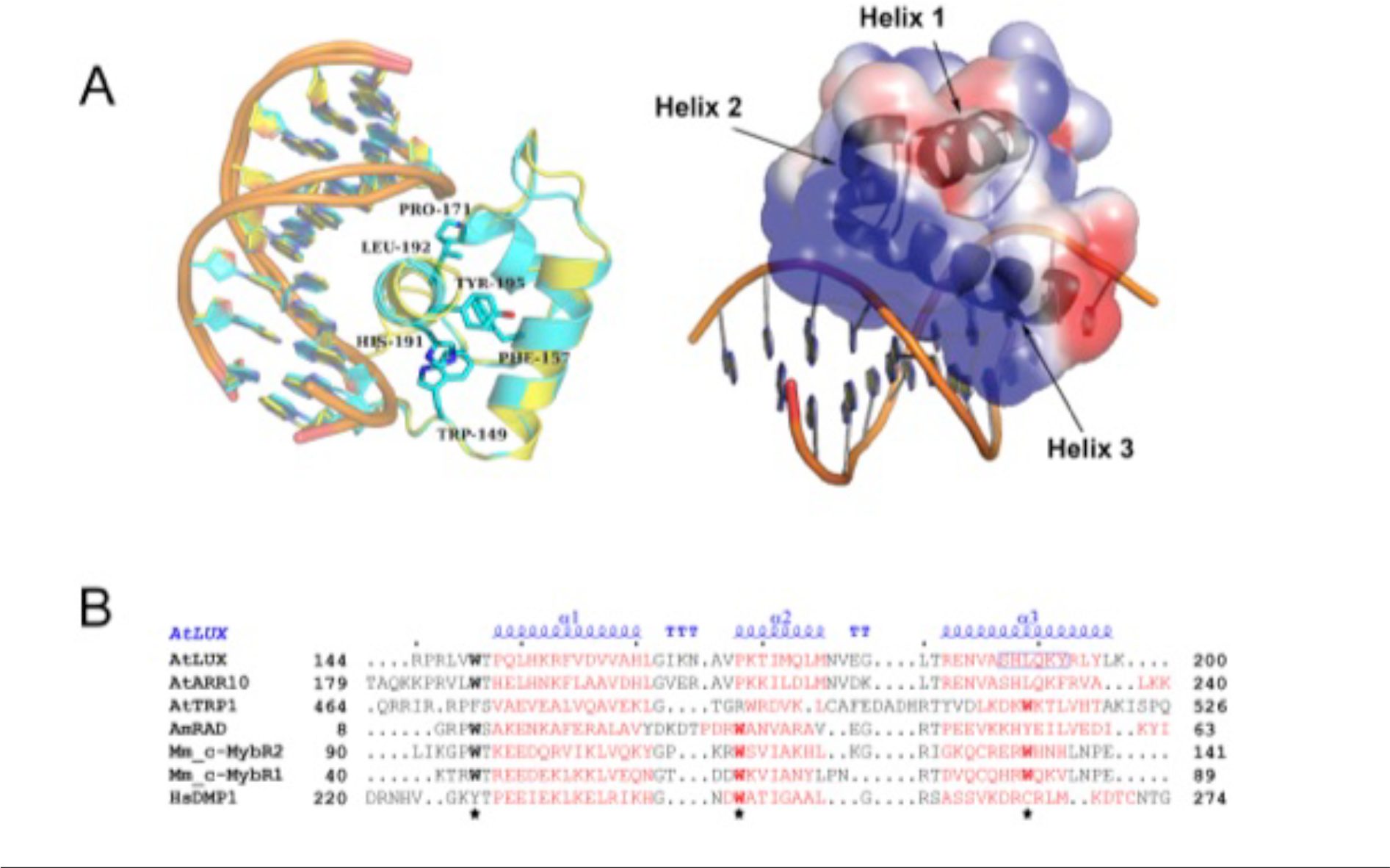
Structure of LUX^MYB^ in complex with DNA. **A.** *Left*, overlay of LUX^MYB^ structures in cyan (PDB 5LXU) and yellow (PDB 6QEC) with the hydrophobic core residues displayed as sticks and colored by atom with carbons in cyan (only 5LXU side chains are shown for clarity). The DNA sequences 5’-TAGATACGCA (cyan carbons) and 5’-TATATTCGAA (yellow carbons) are shown as sticks. *Right*, electrostatic surface representation with electropositive to electronegative surfaces colored from blue to red with helices indicated by arrows. **B.** Structure-based sequence alignment of representative MYB domains; the three regularly spaced hydrophobic residues characteristic of MYB domains are indicated with a star. Depicted in red are the α-helices derived from each corresponding structure. The secondary structure annotation of LUX^MYB^ is depicted in blue on top of the aligned sequences (α, alpha helices; TT, strict β-turn; TTT, strict α-turn).

Helix 2 and 3 form a helix-turn-helix motif, constituting an electropositive groove for the negatively charged DNA and acting as the primary interface with the LBS. Residues from helix 3 account for the majority of the direct base readout and also contribute sugar-phosphate backbone interactions between the protein and DNA (Figure 4A). While no residues in helix 1 directly interact with the DNA, Arg146, part of the unstructured N-terminal extension, intercalates into the minor groove and interacts largely via van der Waal’s forces and a water-mediated hydrogen bonding network with adenine and thymine/guanine of the bound DNA (T<u>**AG**</u>ATACGA and T<u>**AT**</u>ATTCGAA) (Figure 4B). Interestingly, the Arg146 residue adopts different conformations in the two structures with different hydrogen bonding networks, suggesting plasticity in how Arg146 binds DNA.

**Figure 4.**
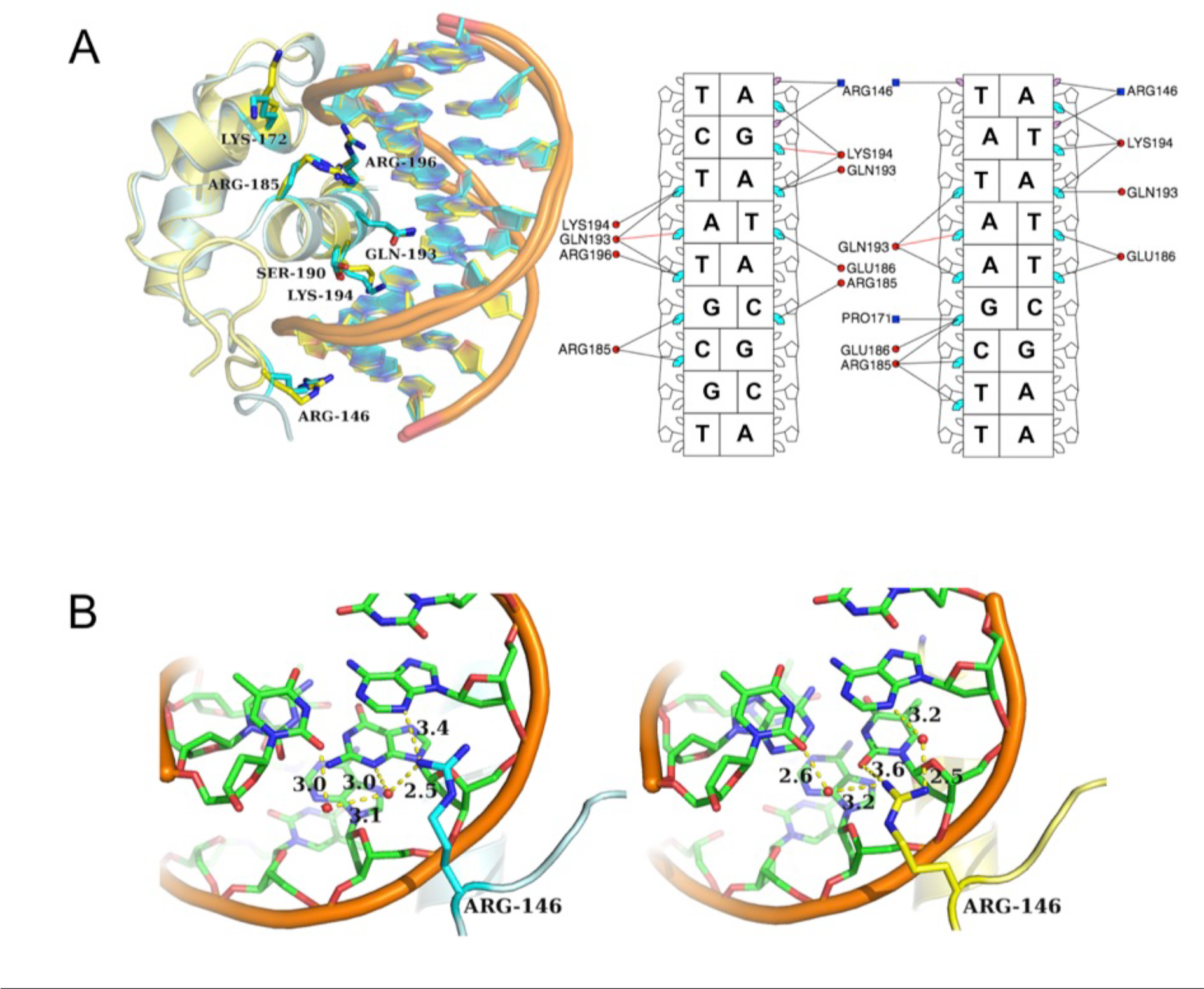
LUX^MYB^-DNA interactions. **A.** *Left*, overlay of LUX^MYB^-DNA structures. Amino acid residues interacting with the DNA are shown as sticks and colored by atom with carbons in cyan (5’-TAGATACGCA) or yellow (5’-TATATTCGAA), DNA is shown as a cartoon. *Right*, simplified schematic from DNAproDB (53) of amino acids important for DNA-binding and base read-out with only direct base interactions shown for clarity. The LBS sequences are depicted without the first overhanging base, major groove interactions are shown in cyan, minor groove interactions in purple, protein helices are in red circles and loop residues are in blue squares. **B.** Close-up view of Arg146 interactions with DNA colored by atom with carbons in green. The proteins are colored as per **A**. Hydrogen bonding interactions are shown as dotted lines and distances labelled. Water molecules are shown as red spheres. Arg146 adopts different conformations in the two structures.

As Arg146 seems to act as a general “clamp” targeting the DNA minor groove, this residue was targeted for mutagenesis. The R146A mutation in both the LUX^MYB^ and full-length constructs was assayed for DNA-binding by EMSA. As predicted, the R146A mutation decreased the binding affinity for both LUX^MYB^ and the full-length protein, albeit with a greater effect depending on the DNA sequence (Table 1).

### Tuning EC activity *in planta*

Based on the *in vitro* results presented above, the ability to alter EC activity through changes in the DNA-binding affinity of LUX was investigated. We hypothesised that decreasing LUX DNA-binding affinity would result in less active EC target gene repression and a warm temperature phenotype intermediate between wild type and the *lux* null mutant. To test this, we examined hypocotyl length under short day conditions at 22°C and 27°C and flowering time under long day conditions at 22°C. The null mutant, *lux-4*, in the Arabidopsis Col-0 background was transformed with either wild type *LUX* or *LUX*^*R146A*^ under the control of its native promoter and hypocotyl length was measured for three independent homozygous lines for each genotype. Transformation with the *pLUX::LUX* construct resulted in complementation based on hypocotyl length (Figure 5 A-C) and on flowering time, measured as the number of rosette leaves at time of bolting, (Figure 5 D, E). This phenotype was recapitulated in subsequent generations.

**Figure 5.**
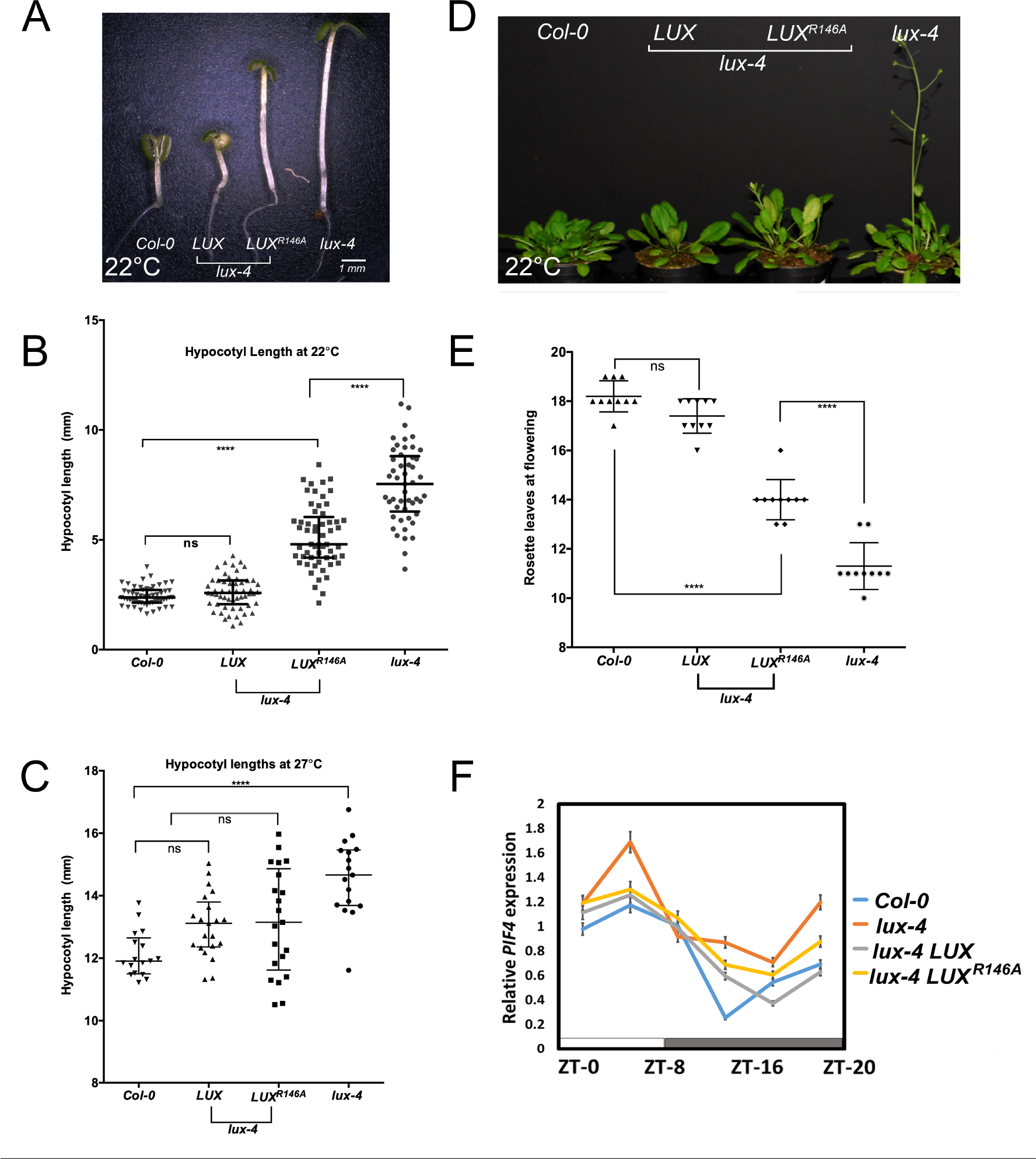
Hypocotyl and flowering phenotypes for *Col-0, lux-4* and transformed lines. **A.** Representative hypocotyls from 7-days-old seedlings grown at 22°. **B.** Hypocotyl length measurements from seedlings grown at 22°C. One-way anova test was performed (ns, not significant, ****, P<0.001). The error bars represent the median value with interquartile ranges. **C**. Hypocotyl length measurements from seedlings grown at 27°C. **D.** Representative images of plants grown at 22°C on soil. Genotypes are indicated. **E**. Number of rosette leaves at time of bolting from indicated genotypes. Error bars represent the mean with standard deviation. **F.** *PIF4* expression over a 24-hour period for seedlings grown at 22°C for the different genotypes. Day and night are indicated as a bar below the graph. Error bars are indicated.

In contrast, *LUX*^*R146A*^ was not able to completely rescue the *lux-4* mutation and an intermediate phenotype between wild type and *lux-4* was observed for both hypocotyl length and flowering time (Figure 5 A-E). Temperature responsive growth was still observed in *LUX*^*R146A*^ but was more attenuated than in wild type or *lux-4* complemented with the wild type gene (Figure 5C). Taken together, these data show that reduction in the binding affinity by the LUX^R146A^ mutation, as demonstrated *in vitro*, has a predictable *in vivo* effect of titrating activity of the entire EC.

### *PIF4* expression is misregulated in *LUX*^*R146A*^

*PIF4* is a master regulator of cell elongation, hypocotyl length and integrates light and temperature signals for plant growth. The *PIF4* promoter contains a consensus LBS sequence (GAT(A/T)CG) and *PIF4* is a target of the EC based on ChIP-seq and RNA-seq experiments (12, 14, 15). Therefore, the expression levels of *PIF4* were examined by RT-qPCR at 4-hour intervals over a 24hr period in the different genotypes (Figure 5F). In wild type Arabidopsis, *PIF4* expression is under circadian control and oscillates over a 24-hr period, with a strong repression at ZT12 when EC expression peaks, as shown in Figure 5F. *PIF4* expression increases as EC expression decreases, peaking during the day. In the *lux-4* mutant, *PIF4* expression levels are elevated with respect to WT and the sharp decrease in *PIF4* expression at dusk (ZT8-ZT12) is attenuated. The *lux-4* mutant is complemented by *pLUX::LUX* based on hypocotyl length and flowering time, however some misregulation of PIF4 is still observed as wild type levels of PIF4 repression are not fully restored. This misregulation is more evident in *lux-4 pLUX::LUX*^*R146A*^ and is consistent with the observed intermediate phenotype between wild type and *lux-4*. Expression of *PIF4* was elevated with respect to wild type in *LUX*^*R146A*^ and did not exhibit the sharp decrease in expression between ZT8 and ZT12, similar to the *lux-4* mutant. Overall, *PIF4* expression was elevated in the *LUX*^*R146A*^ lines compared to wild type, but lower than in the *lux-4* mutant suggesting that LUX^R146A^ triggers a misregulation of *PIF4*. It is likely that the observed phenotypes are due at least in part to changes in *PIF4* expression in the different mutant lines.

## Discussion

The EC plays an important role in the circadian clock, functioning to connect the evening and morning loops by forming a key circuit in the plant circadian system (33). In addition, the EC is ideally positioned to act as a hub for integrating environmental cues and relaying this information directly to growth and developmental pathways through direct effects on target genes including *PIF4*, a master regulator of thermosensory growth, plant immunity and reproductive development (26, 34, 35). Using the EC to tune temperature sensitive growth is an attractive goal requiring an understanding of the molecular basis of EC formation and activity. However, this was poorly defined. To address this deficit we sought to provide a molecular model of LUX, ELF3 and ELF4 interactions in EC formation and to define the residues critical for DNA-binding.

Regulation of target genes requires the full EC as suggested by the similar phenotypes of *lux-4*, *elf3-1* and *elf4* mutants (17, 36–38), however only LUX possesses a DNA binding domain. Indeed, LUX binding to DNA *in vivo* was observed in the *elf3-1* background, demonstrating that complex formation is not required for LUX DNA-binding (15) but is required for regulation. LUX and ELF3 have been shown to interact in yeast two-hybrid assays and *in vivo* (12). Here, we demonstrate that *in vitro*, ELF3 is able to impede LUX-DNA interactions, at least for certain LBS sequences, suggesting the formation of a complex that is not competent to bind DNA. This type of sequestering role of ELF3 has been observed for ELF3-PIF4 interactions and may be a general function of ELF3 in other protein complexes (39). We further demonstrate that ELF4 modulates the activity of ELF3 and restores DNA-binding competency to the LUX-ELF3-ELF4 complex (EC), highlighting its crucial role in the EC. Indeed, previous modelling studies of the contributions to EC activity suggested that *ELF4* transcript levels are as powerful a predictor of EC target gene repression as using the full EC (*LUX*, *ELF3* and *ELF4* transcript levels) and more predictive than *ELF3* alone (15). Thus, only with all three components, LUX, ELF3 and ELF4, do both DNA-binding and target gene regulation occur.

Based on the *in vitro* and structural studies presented here, LUX provides the specificity and affinity necessary to target the entire EC to its cognate binding sites. The MYB domain is able to perform direct base read-out of the core LBS. The plant specific signature sequence, SH(A/L)QK(F/<u>Y</u>) of helix 3, provides the majority of direct interactions via the major groove. In addition, an N-terminal arginine, Arg146, part of a flexible extension, is important for intercalation into the minor groove and acts as a DNA clamp. Arginine residues in flexible extensions are found in many other structurally diverse TFs including homeodomain TFs and MADS TF family members (40, 41). While these arginine residues are likely important for DNA shape readout by intercalating into the minor groove (42), they are often not required for direct base-readout and may offer a general way to decrease DNA binding affinity while maintaining specificity for other TF families, although this remains to be tested.

Arg146 of LUX was mutated to alanine in order to decrease the DNA-binding affinity of the EC for its cognate sites while still retaining specificity. Based on *in vitro* assays, we predicted dampened but not abolished EC activity *in planta.* Indeed, at 22°C an intermediate early flowering phenotype between wild type and *lux-4* was observed for *lux-4* plants transformed with *pLUX::LUX*^*R146A*^. EC activity is greatly reduced at 27°C and this was reflected in the similar phenotypes of wild type and the *lux-4* transformed lines. The R146A mutation resulted in accelerated growth but still retained thermo-response.

*PIF4* is an important direct target of the EC implicated in hypocotyl elongation and thermoresponsive growth and a target of the EC (15, 26, 35). PIF4 expression was higher in the *lux-4 pLUX::LUX*^*R146A*^ plants as compared to wild type and these plants phenocopy mild *PIF4* over-expressors which have an early flowering phenotype and elongated hypocotyls at 22°C (35). The peak of PIF4 expression was similar for wild type and *lux-4 pLUX::LUX*^*R146A*^, however the characteristic strong decrease in PIF4 expression between ZT8 and ZT12, which coincides with maximum EC expression, was less apparent, likely due to the decreased affinity of the R146A mutation for its LBS.

The roles of the EC as a core clock component and as a direct regulator of *PIF4* make it an attractive target for tuning plant development. The EC is present in basal plants such as mosses to higher flowering plants, suggesting an evolutionarily conserved function. Altering EC activity via structure-based design or directed evolution is a potential strategy to modify plant growth and flowering pathways in not only model plants but also crop species. Directed evolution strategies have been used to alter transcription factor properties including changing DNA-binding specificities and increasing/decreasing DNA-binding affinity (43–45). The data presented here demonstrate that rational design strategies targeting amino acids important for LUX binding can be used to tune the activity of the entire EC, resulting in plants with a predictable phenotype, suggesting a viable method for rational engineering of plant development.

## Materials and Methods

### Protein binding microarrays

LUX (LUX FL; TAIR At3g46640.1) and LUX^MYB^ (amino acid residues 139-200wer cloned into the pETM41 vector and recombinant proteins were expressed as described (30, 31). DNA-binding specificities were determined using protein binding microarrays (PBM11), as previously described (30, 31).

### Protein expression

*LUX, LUX*^*MYB*^ and *ELF4* were cloned into the expression vector pESPRIT002 (46, 47), expressed and purified using standard protocols (46, 47). Seleno-methionine (SeMet) derived LUX^MYB^ protein was produced according to standard protocols (48).

Full-length *ELF3* (TAIR At2g25930.1) was cloned into the pACEBac1(49) vector and produced in *Sf*21 insect cells (Invitrogen) using the baculovirus expression system.

### Protein Purification

LUX^FL^, LUX^MYB^ and SeMet LUX^MYB^ proteins were isolated following the same purification protocol. Harvested cells were resuspended in 200 mM CAPS pH 10.5, 500 mM NaCl, 1 mM TCEP, sonicated, centrifuged and the soluble proteins purified by Ni-affinity chromatography. The N-terminal 6xHis tag was cleaved with TEV protease and the protein further purified using a heparin (LUX^MYB^) or Superdex 200 (LUX^FL^) column (GE Healthcare).

For ELF4 protein, harvested cells were resuspended in 20 mM Tris pH 8.0, 500 mM NaCl, 1 mM TCEP. Purification was as per LUX^FL^.

For ELF3 the protein was purified from 8 M Urea using a Ni-Sepharose High-Performance resin column. The protein was refolded in the presence of LUX and ELF4 through step-wise dialysis from 8M to 0M urea.

### Protein crystallisation and data collection

Protein-DNA complexes were crystallised using the hanging drop method as previously described (50).

Diffraction data were collected at 100K at the European Synchrotron Radiation Facility (ESRF), Grenoble France. Data collection and refinement statistics are given in SI Table 1. The structures are deposited under PDB codes 5LXU and 6QEC.

### Sequence Alignments

Structure-based sequence alignments were performed using the server PROMALS3D (51). All structures were aligned with TM-align using the default parameters (52).

### Electrophoretic Mobility Shift Assays (EMSAs)

DNA was Cy5 labelled for visualisation and used at a final concentration of 10 nM for PAGE and 30 nM for agarose gels. Protein and DNA were incubated at room temperature in binding buffer (10 mM Tris pH 7.0, 50 mM NaCl, 1 mM MgCl2, 1 mM TCEP, 3% glycerol, 28 ng/μL herring sperm DNA, 20 μg/mL BSA, 2.5% CHAPS and 1.25 mM spermidine) and protein-DNA complexes run on a 8% polyacrylamide gel using TBE buffer 0.5x in non-denaturing conditions at 4°C.

Protein concentration was varied from 0 nM to 1000 nM for LUX^MYB^ and LUX^FL^ experiments. For LUX-ELF3 and LUX-ELF3-ELF4 experiments, all tested complexes were reconstituted by mixing the proteins of interest in 6 M urea followed by a step-wise dialysis to 0 M urea. LUX and ELF4 concentrations were constant while the ELF3 concentration was varied from 220 nM to 2.2 μM. Proteins and DNA were incubated at room temperature and protein-DNA complexes run on a 2% agarose gel using TBE buffer 0.5x in non-denaturing conditions at 4°C.

### Plant material and cultivation conditions

For qPCR and hypocotyl measurements material was collected from 7-days old seedlings grown in FitoClima D1200 (Aralab) growth chambers, at 22°C (SD, 8h light/16h dark). Hypocotyl measurements were performed on the T2 generation of plants with 15-25 plants for each independent line. For flowering phenotype analysis, primary transformants were selected for the transgene and sown on soil and transferred to LD conditions after stratification (4°C, 3 days). Flowering time was determined in randomly distributed plants according to number of rosette leaves at the time of bolting (10 plants for wild type, *lux-4*, *lux-4* p*LUX*::*LUX*^*R146A*^ and *lux-4 pLUX::LUX*).

### Plasmid construction and generation of transgenic plants

For the *lux-4* p*LUX*::*LUX*^*R146A*^, *lux-4 pLUX::LUX* constructs, a ~800bp upstream fragment of *LUX* was PCR-amplified from genomic DNA. Full length CDS of *LUX* and *LUX*^*R146A*^ were PCR-amplified from the pESPRIT002 expression vector containing the respective CDS with an N-terminal FLAG tag added (SI Figure 2). NEBuilder^®^ HiFi DNA Assembly Kit (E2621S, NEB) was used for assembly with the vector backbone (pFP101 containing the *At2S3* promoter driven GFP for selection of transformants)(57). For a list of primers see SI Table 2. Transgenic plants were generated using the floral dip method (58).

### RNA isolation and quantitative PCR

Plants were grown under short day conditions (8L:16D) and samples were harvested in intervals of 4 hours. 8-10 seedlings were harvested for each line at each time point. Total RNA was extracted using RNeasy Plant mini kit (Qiagen) according to manufacturer’s instructions. Total RNA (1µg) was treated with DNaseI (Roche) qRT-PCR was done using iTaq^®^ Universal SYBR^®^ Green One-Step Kit from Bio-Rad following manufacturer’s protocol. For the list of primers see **SI Table 2**. Expression of *PIF4* in different plant lines was determined through qRT-PCR with *PP2A* used as a control. qRT-PCR measurements were performed with a Bio-Rad CFX connect Real-Time system. Quantification was performed with the relative –ΔCt method, using *PP2A* for normalization. All quantification and statistical analysis were performed using CFX Maestro^TM^ software (Bio-Rad).

Details for Materials and Methods are given in accompanying SI.

## Acknowledgments

We would like to acknowledge Darren Hart and Philippe Mas for the pESPRIT002 vector and the ESRF beamline staff of ID29, ID23-2 and the EMBL HTX Lab Funding was from the European Community’s Seventh Framework Programme (FP7/2007-2013) under BioStruct-X (grant agreement N°283570), the Raman Chaprak Fellowship (to AN), CEA Irtelis program (AN, CZ), ATIP-Avenir (CZ), GRAL (ANR-10-LABX-49-01) (CZ, VH, FP) and the Spanish MINECO grant BIO2017-86651-P (AEI/FEDER) to JMF-Z

## Supplemental Information

**SI Table 1.**
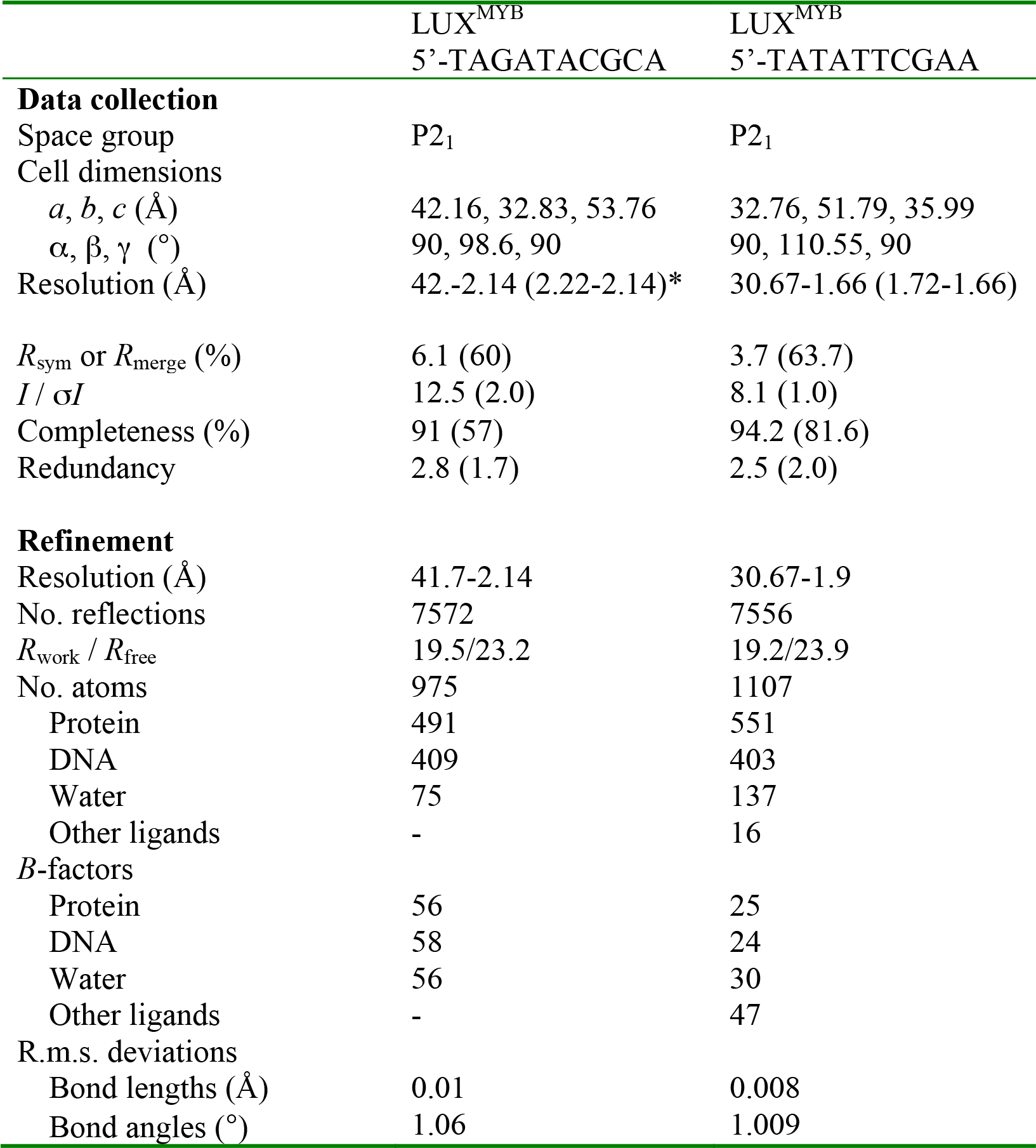
Data collection and refinement statistics

|  | LUX<sup>MYB</sup><br>5'-TAGATACGCA | LUX<sup>MYB</sup><br>5'-TATATTCGAA |
| --- | --- | --- |
| **Data collection** |  |  |
| Space group | P2<sub>1</sub> | P2<sub>1</sub> |
| Cell dimensions |  |  |
| *a*, *b*, *c* (Å) | 42.16, 32.83, 53.76 | 32.76, 51.79, 35.99 |
| $\alpha$, $\beta$, $\gamma$ (°) | 90, 98.6, 90 | 90, 110.55, 90 |
| Resolution (Å) | 42.-2.14 (2.22-2.14)* | 30.67-1.66 (1.72-1.66) |
| *R*<sub>sym</sub> or *R*<sub>merge</sub> (%) | 6.1 (60) | 3.7 (63.7) |
| *I* / $\sigma I$ | 12.5 (2.0) | 8.1 (1.0) |
| Completeness (%) | 91 (57) | 94.2 (81.6) |
| Redundancy | 2.8 (1.7) | 2.5 (2.0) |
| **Refinement** |  |  |
| Resolution (Å) | 41.7-2.14 | 30.67-1.9 |
| No. reflections | 7572 | 7556 |
| *R*<sub>work</sub> / *R*<sub>free</sub> | 19.5/23.2 | 19.2/23.9 |
| No. atoms | 975 | 1107 |
| Protein | 491 | 551 |
| DNA | 409 | 403 |
| Water | 75 | 137 |
| Other ligands | - | 16 |
| *B*-factors |  |  |
| Protein | 56 | 25 |
| DNA | 58 | 24 |
| Water | 56 | 30 |
| Other ligands | - | 47 |
| R.m.s. deviations |  |  |
| Bond lengths (Å) | 0.01 | 0.008 |
| Bond angles (°) | 1.06 | 1.009 |

**SI Table 2.**
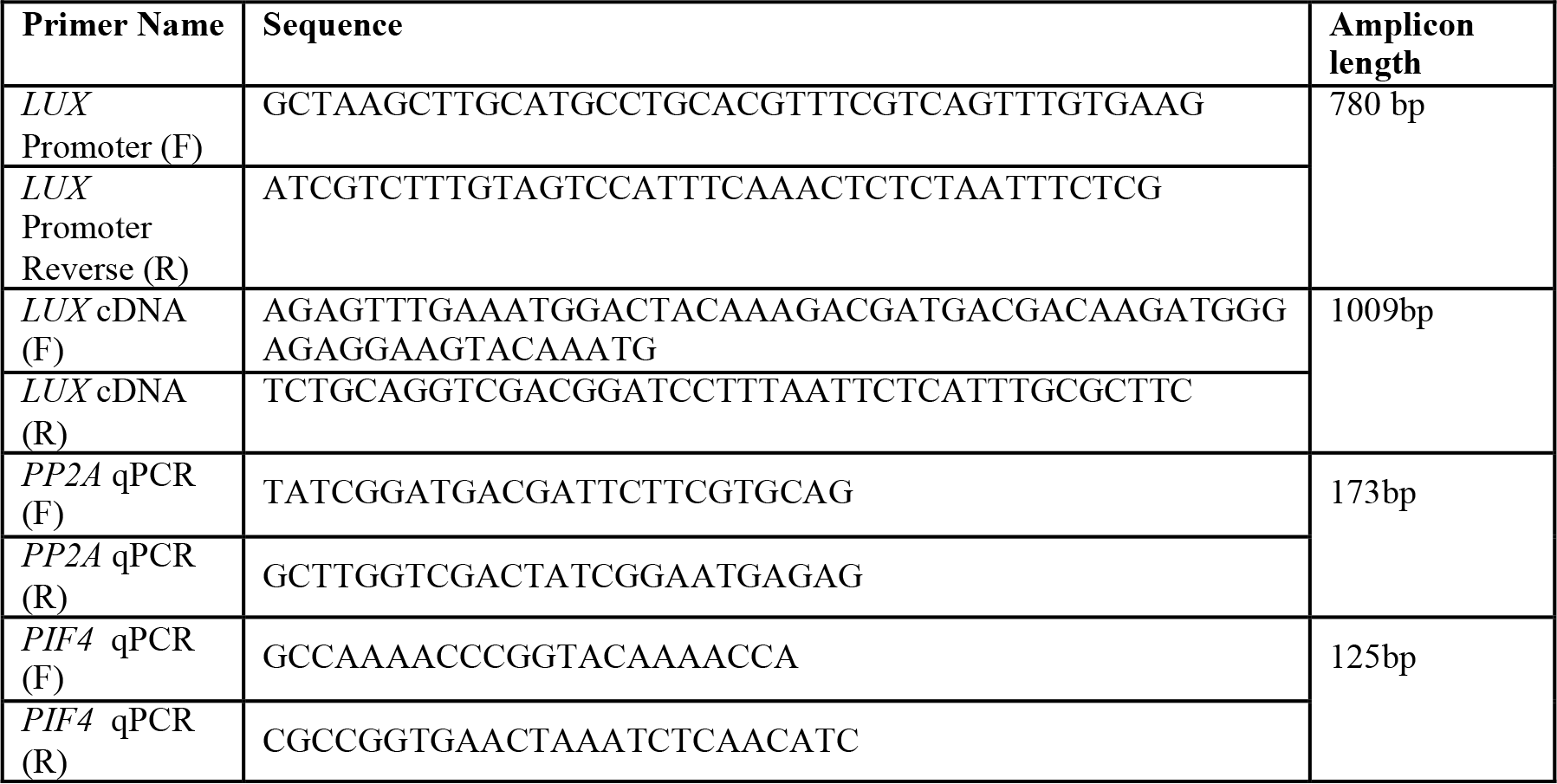
Primers used for design of plant expression vector and qRT-PCR (F=forward primer, R=reverse primer)

**SI Figure 1.**
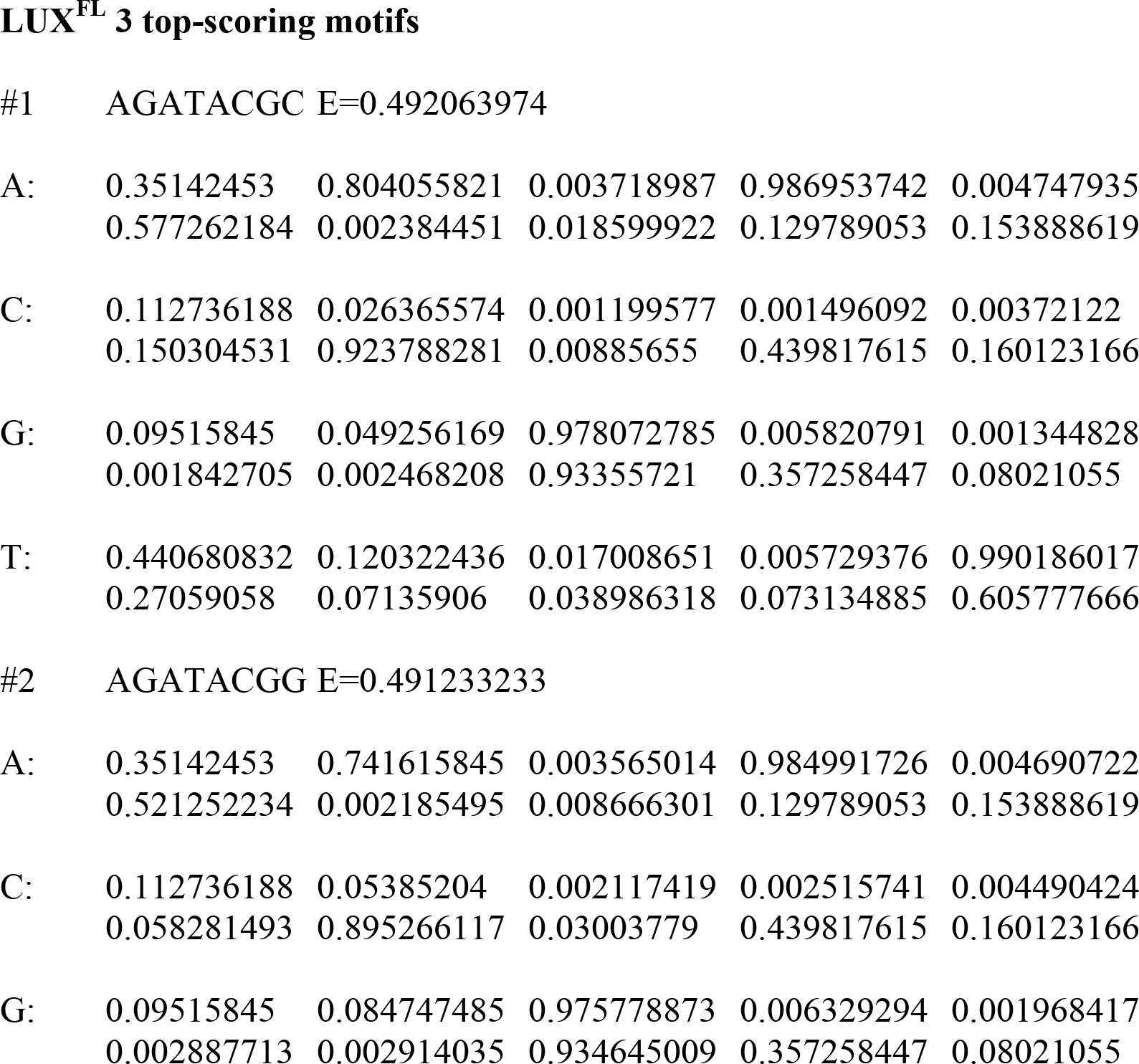
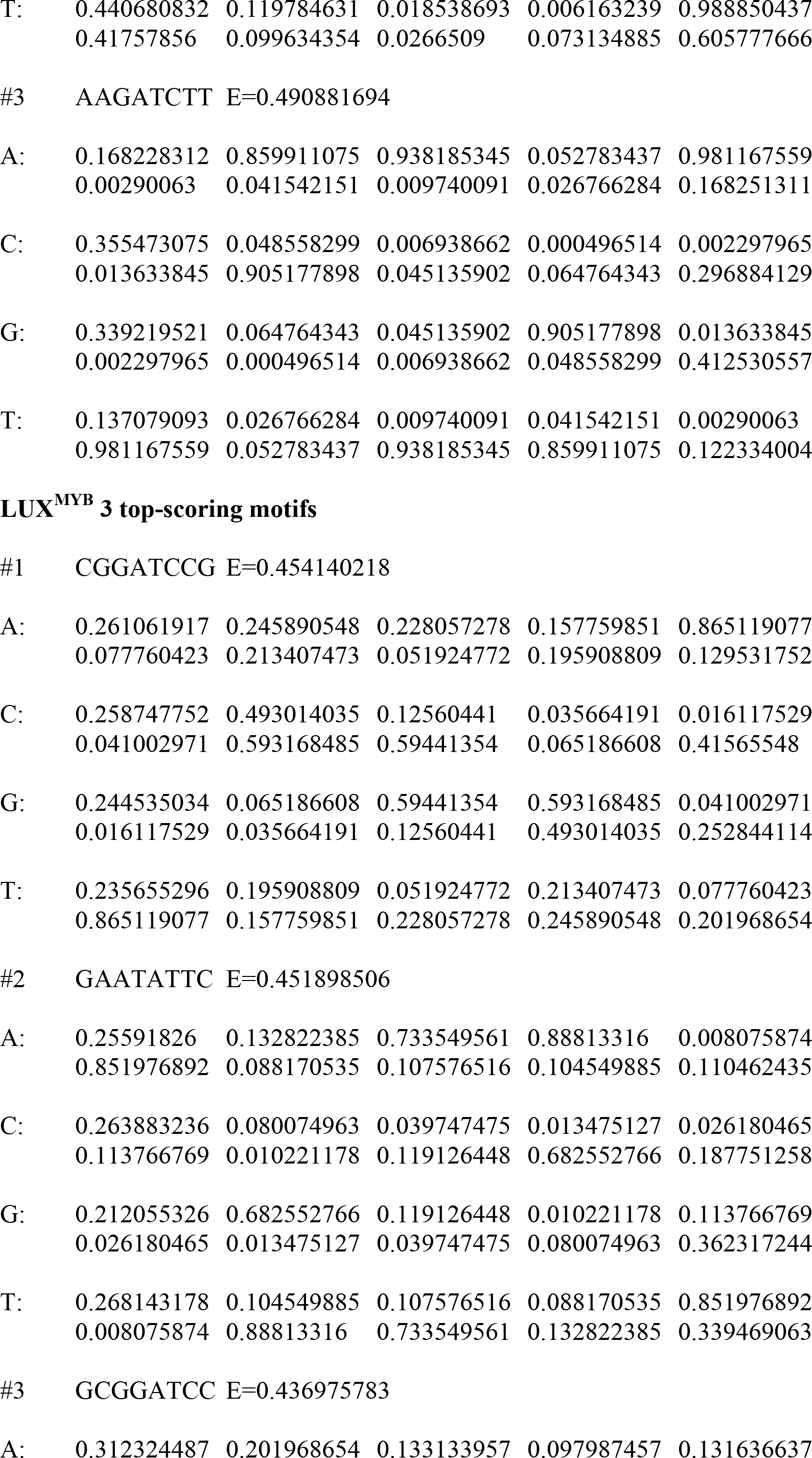
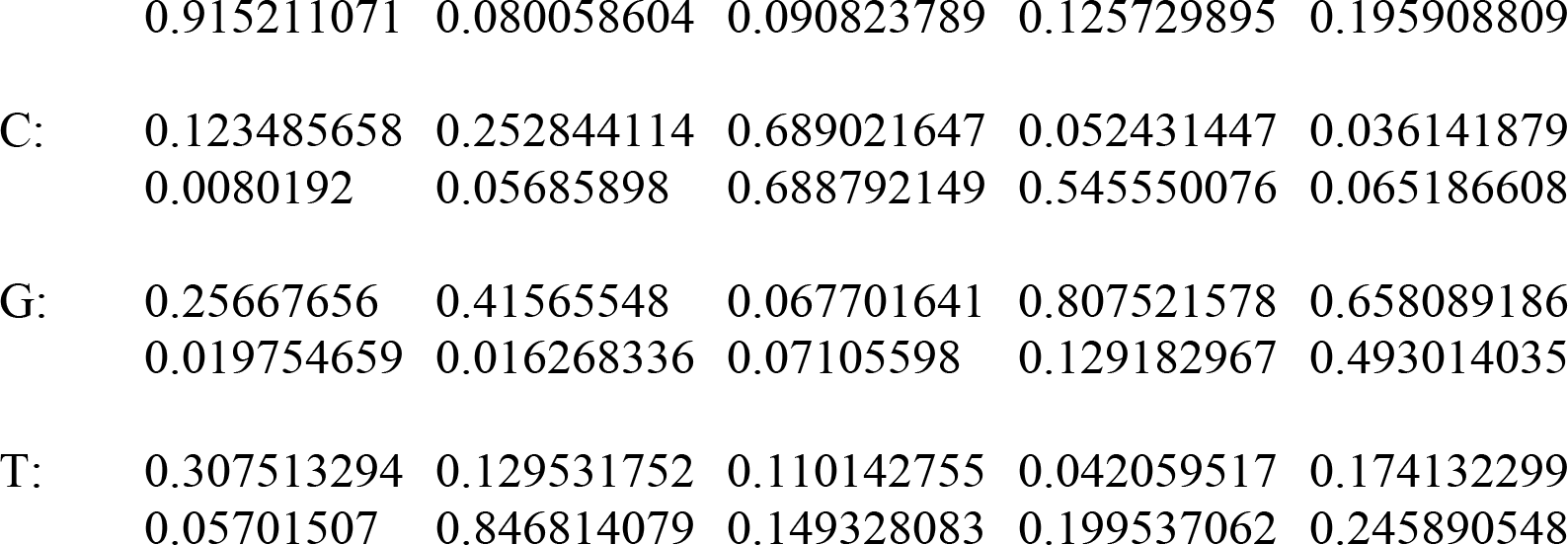
LUX^FL^ and LUX^MYB^ 3 top-scoring motifs and position weight matrices LUX^FL^ 3 top-scoring motifs

**SI Figure 2.**
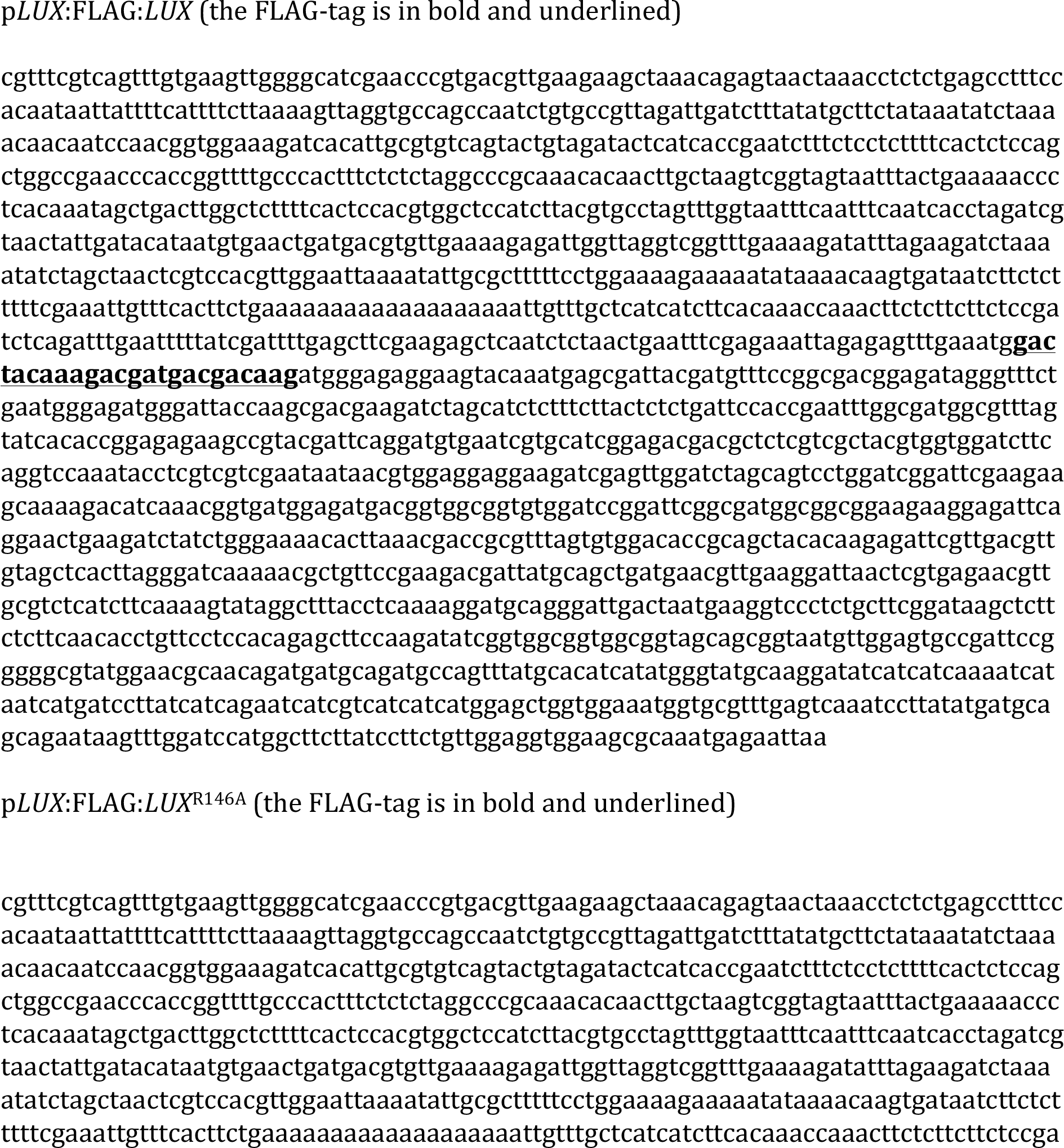
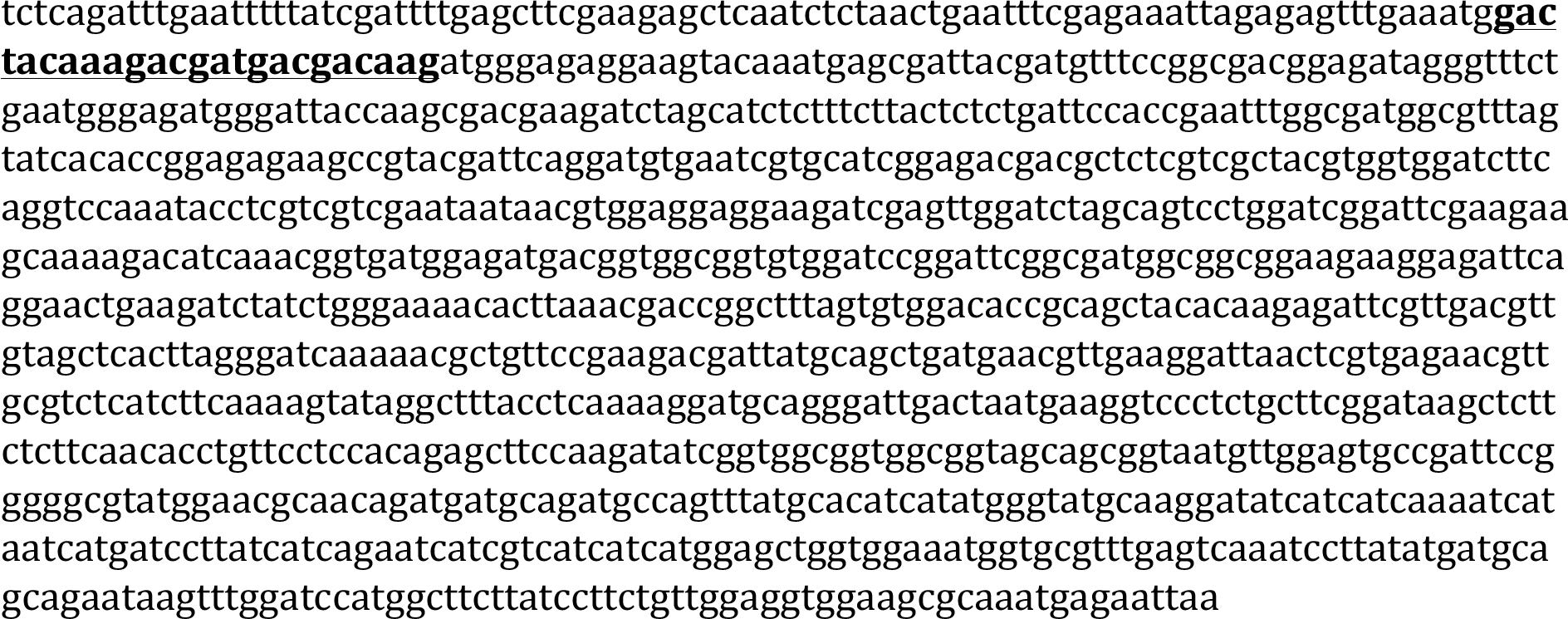
DNA sequences used for plasmid constructions for transformation in plants.

## SI Materials and Methods

### Protein binding microarrays

Translational fusions of LUX (LUX FL; TAIR At3g46640.1) and LUX^MYB^ (amino acid residues 139-200) to Maltose Binding Protein were obtained by cloning their corresponding cDNAs into the pETM41 vector (EMBL) using restriction sites NcoI and NotI. Clones were sequence verified and the plasmids introduced into *Escherichia coli* BL21. Recombinant proteins were expressed as described (30, 31). DNA-binding specificities were determined using protein binding microarrays (PBM11), incubated with soluble protein extracts obtained from 25 mL induced *E. coli* cultures. Incubation of protein extracts and antibodies, washing, scanning, quantification and analysis were performed as described (30, 31). The top 3 highest scoring motifs are given in SI Figure 1.

### Protein expression

The full-length LUX was cloned into the expression vector pESPRIT002(46, 47) using the AatII and NotI restriction sites. A second construct LUX^MYB^ (amino acid residues 139-200) was generated following the same strategy. Both constructs contained a TEV protease cleavable N-terminal 6x-His tag. LUX^FL^ and LUX^MYB^ constructs were overproduced in *Escherichia coli* (*E.coli*) Rosetta2 (DE3) pLysS cells. Cells were grown at 37°C in Luria-Bertani (LB) culture medium supplemented with chloramphenicol (37 mg/mL) and kanamycin (50 mg/mL), until an OD_600_ of 0.7-0.8. The temperature was then reduced to 20°C and protein expression induced by addition of 1 mM isopropyl-β-D-thiogalactopyranoside (IPTG); expression was continued overnight (~16 h) and the cells were harvested by centrifugation for 30 min. at 6000 rpm at 4°C.

Seleno-methionine (SeMet) derived LUX^MYB^ protein was produced according to standard protocols (48). Briefly, SeMet LUX^MYB^ was produced in M9 minimal medium using a non-auxotrophic strain (Rosetta2 (DE3) pLysS cells). Overnight grown LB precultures were spun down, washed with M9 medium and used to inoculate M9 cultures supplemented with chloramphenicol (37 mg/mL), kanamycin (50 mg/mL), MgSO_4_ (2 mM), glucose (0.4% w/v) and MgCl_2_ (0.1 mM). Cells were grown at 37°C until an OD_600_ of 1.2 was reached. Amino acids (100 mg/L lysine, threonine and phenylalanine; 50 mg/L leucine, valine, isoleucine and L-seleno-methionine) were added and the temperature reduced to 20°C. After 15 minutes, 0.5 mM IPTG was added to induce protein expression and the culture grown overnight. Cells were harvested as for the native protein.

*ELF4* full-length coding sequence (TAIR At2g40080.1) was cloned into the expression vector pESPRIT002(46, 47) following the same strategy as for LUX^FL^. The resulting construct, containing a TEV cleavable N-terminal 6x-His tag, was overproduced in *Escherichia coli* (*E.coli*) BL21 CodonPlus RIL cells under the same conditions as LUX^FL^.

Full-length *ELF3* (TAIR At2g25930.1) was cloned into the pACEBac1(49) vector via SalI and NotI restriction sites and inserting a TEV protease cleavable 6x-His sequence at both N- and C-terminal ends of the gene. ELF3 protein was produced in *Sf*21 insect cells (Invitrogen) using the baculovirus expression system. Briefly, the generated construct was transformed into chemically competent DH10 EmBacY cells (harbouring the baculoviral EmBacY genome) (54, 55). Positive clones were identified by blue/white screening in the presence of IPTG and BluoGal and used for downstream bacmid isolation. *Sf*21 insect cells were transfected at a density of 0.3×10^6^ cells/mL in a 6-well plate format. Primary baculovirus stock (V_0_) was harvested 60 h after transfection and used for infecting 25 mL new *Sf*21 cell cultures yielding V_1_ stock. V_1_ baculovirus stock was collected 48h after cell proliferation arrest, stored at 4°C and used to launch ELF3 expressions (V_2_) (500 mL cell cultures at 1.0×10^6^ cells/mL with 0.1% (v/v) V_1_ virus). Amplification of the virus and protein expression were followed by monitoring YFP (Yellow Fluorescent Protein) expression from the viral backbone and cells harvested (2000g, 15 min, 4°C) when reaching a fluorescence signal plateau. All experiments were performed at 27°C.

### Protein Purification

LUX^FL^, LUX^MYB^ and SeMet LUX^MYB^ proteins were isolated following the same purification protocol. Harvested cells were resuspended in buffer A (200 mM CAPS pH 10.5, 500 mM NaCl, 1 mM TCEP) to which benzonase and protease inhibitors were added. Cells were disrupted by sonication, followed by centrifugation for 40 min. at 25000 rpm and 4°C. Cell lysates were applied onto a 1 mL Ni-Sepharose High-Performance resin (GE-Healthcare) column, pre-equilibrated with Buffer A. The column was then washed with 15 CV of wash buffer (buffer A + 10 mM imidazole) and the protein subsequently eluted with buffer B (buffer A + 200 mM imidazole). Fractions of interest were pooled and dialysed overnight at 4°C against dialysis buffer (50 mM CAPS pH 9.7, 500 mM NaCl, 1mM TCEP) in the presence of 2% (w/w) TEV protease, in order to cleave the N-terminal 6xHis tag. The protein samples were then concentrated and buffer exchanged against buffer C (50 mM CAPS pH 9.7, 100 mM NaCl, 1 mM TCEP) before being applied onto a 1 mL heparin column (GE-Healthcare) and eluted against a 25 CV salt gradient (buffer D: buffer C + 1 M NaCl). LUX^MYB^ and SeMet LUX^MYB^ fractions of interest were pooled after the heparin column, buffer exchanged with buffer C and concentrated to 5-10 mg/mL, for crystallisation studies. As LUX^FL^ exhibited poor binding to the heparin column, the flow through was collected, pooled, concentrated and passed over a size exclusion Superdex 200 10/300 GL column (GE-Healthcare), pre-equilibrated with buffer C, as a final purification step. LUX^FL^ was concentrated to 14 mg/mL.

For ELF4 protein, harvested cells were resuspended in 20 mM Tris pH 8.0, 500 mM NaCl, 1 mM TCEP (buffer E) to which benzonase and protease inhibitors were added. Cells were lysed by sonication, cell debris removed via centrifugation for 40 min. at 18000 rpm and 4°C, and the supernatant applied onto a 1 mL Ni-Sepharose High-Performance resin column, pre-equilibrated with Buffer E. The column was washed with 25 CV of wash buffer (buffer E + 20 mM imidazole) and the protein eluted with buffer F (buffer E + 200 mM imidazole). Fractions of interest were pooled and dialysed overnight at 4°C against buffer G (20 mM Tris pH 8.0, 300 mM NaCl, 1 mM TCEP) and in the presence of 2% (w/w) TEV protease, in order to cleave the N-terminal 6xHis tag. The protein sample was then passed onto a second Ni-Sepharose column (150 μL resin) in order to deplete 6xHis tagged TEV protease and subsequently applied to a size exclusion Superdex 200 Hi-Load 16/60 column (GE-Healthcare), pre-equilibrated with buffer H (20 mM Tris pH 8.0, 100 mM NaCl, 1 mM TCEP). Protein fractions were pooled and concentrated to 10-16 mg/mL.

For ELF3 the cell pellet was resuspended in buffer I (8 M Urea + 1 mM TCEP) and lysed via four freeze/thaw cycles in liquid nitrogen and an ice/water bath. The cell lysate was then centrifuged for 1h at 18000 rpm and 4°C and the supernatant applied to a 0.5 mL Ni-Sepharose High-Performance resin column, pre-equilibrated in buffer I. The column was washed with 15 CV of wash buffer (buffer I + 20 mM imidazole) and the protein subsequently eluted with elution buffer (buffer I + 200 mM imidazole). Fractions of interest were pooled and dialysed overnight at 4°C against 7 M Urea + 1 mM TCEP, followed by a 4h dialysis against 6 M Urea + 1 mM TCEP. The protein sample was then concentrated up to 1-1.5 mg/mL.

### Protein crystallisation and data collection

A protein-DNA complex was prepared for SeMet derivatised LUX^MYB^, using a 1:1.2 protein:DNA molar ratio. The 10-mer dsDNA sequence with a one base overhang on the 5’ and 3’ ends (forward oligo 5’-TAGATACGCA-3’, reverse oligo 5’-ATGCGTATCT-3’) was ordered as single stranded oligonucleotides (Eurofins). Equimolar concentrations of the two oligonucleotides were mixed, heated to 95°C for 5 min., annealed and used without further purification. Crystallisation experiments were carried out by the vapour diffusion method at 293K, using sitting-drops with a 1:1 ratio of protein-DNA complex:precipitant with a protein concentration of ~6 mg/mL. Suitable well-diffracting crystals were grown after 2-4 days in 0.1 M BisTris Propane, pH 6.5, 20% PEG 3350 and 0.2 M sodium malonate. Crystals grew as needles to dimensions of 200×50×50 μm and were harvested and flash frozen in liquid nitrogen. For the native LUX^MYB^ structure with the DNA sequence (5’-TATATTCGAA-3’, reverse oligo 5’-ATTCGAATAT-3’) crystallisation conditions were the same as above. Crystallisation was performed by the EMBL High Throughput Crystallisation Facility (HTX).

Diffraction data were collected at 100K, on beamline ID29 at the European Synchrotron Radiation Facility (ESRF), Grenoble France. A data set was collected to 2.1Å at the peak absorbance of selenium (0.97886 Å) (50). Indexing was performed using EDNA(56) and the default optimized oscillation range and collection parameters used for data collection. The data set was processed and scaled using the programs XDS and XSCALE.(57) The data were phased using SHELX (58). All refinements were performed using BUSTER (59). Final Ramachandran statistics were 100% preferred region for all residues. The structure is deposited under PDB code 5LXU. For the LUX^MYB^ structure with the DNA sequence (5’-TATATTCGAA-3’, reverse oligo 5’-ATTCGAATAT-3’) data were collected on beamline ID23-1 of the ESRF at 0.976 Å. Indexing was performed using EDNA(56) and the default optimized oscillation range and collection parameters used for data collection. The data set was processed and scaled using the programs XDS and XSCALE(57). Due to anisotropy of the data, the Global Phasing Limited STARANISO server was used for further data reduction and subsequent refinements (http://staraniso.globalphasing.org/cgi-bin/staraniso.cgi). All refinements were performed with Phenix. Final Ramachandran statistics were 100% preferred region for all residues. The structure is deposited under PDB code 6QEC. Data collection and refinement statistics are given in **Table 2**.

### Sequence Alignments

Structure-based sequence alignments were performed using the server PROMALS3D (51). The three-dimensional structures of AtLUX (PDB 5LXU), AtARR10 (*Arabidopsis thaliana* Type-B Response Regulator 10, PBD 1IRZ), AmRAD (*Antirrhinum majus* RADIALIS, PDB 2CJJ), AtTPR1 (*Arabidopsis thaliana* Telomeric-Repeat-binding Protein 1, PDB 2AJE), HsDMP1 (*Homo sapiens* cyclin D-binding Myb-like TF 1, PDB 2LLK) and Mm_c-MybR1 and R2 (*Mus musculus* c-Myb Repeat 1 and 2, PBD 1GUU and 1GV5) were aligned with TM-align using the default parameters (52).

### Electrophoretic Mobility Shift Assays (EMSAs)

A 36bp DNA oligonucleotide (5′-ATGATGTCTTCTCAA<u>**GATTCGA**</u>TAAAAATGGTGTTG-3’) from the *PRR9* promoter containing a LUX DNA-binding site (in bold and underlined) was used for EMSAs, and the core LBS mutated to yield different sequences:

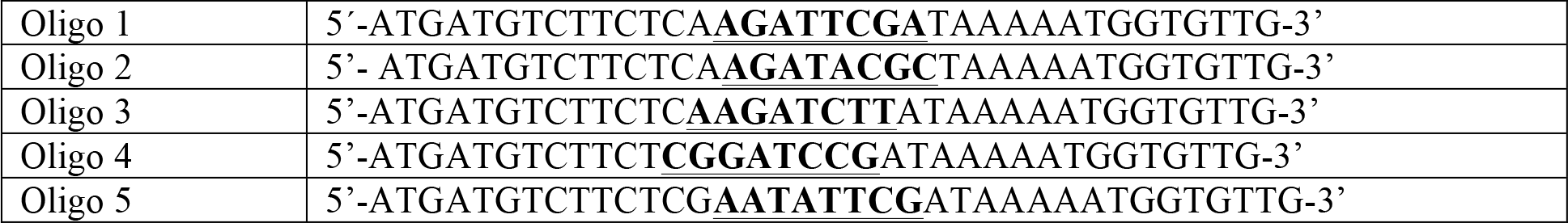

All dsDNA oligonucleotides tested were Cy5 labelled (Eurofins Genomics). Protein concentration was varied from 0 nM to 1000 nM for LUX^MYB^ and LUX^FL^ (0 nM, 2.5 nM, 5.0 nM, 15nM, 30 nM, 60 nM, 120 nM, 250 nM, 500 nM and 1000 nM) using a constant DNA concentration of 10 nM in all reactions. Protein and DNA were incubated at room temperature for 40 min. in binding buffer (10 mM Tris pH 7.0, 50 mM NaCl, 1 mM MgCl2, 1 mM TCEP, 3% glycerol, 28 ng/μL herring sperm DNA (Roche), 20 μg/mL BSA, 2.5% CHAPS and 1.25 mM spermidine) and protein-DNA complexes (LUX^FL^ or LUX^MYB^) run on a 8% polyacrylamide gel using TBE buffer 0.5x in non-denaturing conditions at 4°C.

For LUX-ELF3 and LUX-ELF3-ELF4 experiments, all tested complexes were reconstituted by mixing the proteins of interest in 6 M urea in dialysis buttons, followed by a step-wise dialysis in order to incrementally reduce the urea concentration to 0 M, allowing protein complex refolding (6 M, 5 M, 4 M, 2 M, 1 M urea + 1 mM TCEP in 30 min. steps, and finally 50 mM sodium phosphate pH 7.6 + 100 mM NaCl + 1 mM TCEP for one hour). LUX and ELF4 concentrations were 200 nM and 0 or 1 μM, respectively, while the ELF3 concentration was varied from 220 nM to 2.2 μM. The DNA oligomer used was from the *PRR9* promoter and its concentration maintained constant at 30 nM in all EMSA experiments. Proteins and DNA were incubated at room temperature for 40 min. in binding buffer (10 mM Tris pH 7.0, 1 mM MgCl2, 1 mM TCEP, 6% glycerol, 28 ng/μL herring sperm DNA (Roche), 20 μg/mL BSA, 2.5% CHAPS and 1.25 mM spermidine) and protein-DNA complexes (LUX-ELF3 and LUX-ELF3-ELF4) run on a 2% agarose gel using TBE buffer 0.5x in non-denaturing conditions at 4°C. Gels were scanned using a Chemidoc scanner (Biorad).

### Plant material and cultivation conditions

The *lux-4* mutant allele (background accession Col-0) was provided by Dr. Philip Wigge, Sainsbury Lab, Cambridge University). Seeds of Col-0 and mutants were sterilized in 70% ethanol and sown on 0.5 MS agar medium. For qPCR and hypocotyl measurements material was collected from 7-days old seedlings grown in FitoClima D1200 (Aralab) growth chambers, at 22°C (SD, 8h light/16h dark). Hypocotyl length was measured from images obtained from a flatbed scanner using ImageJ software. Hypocotyl measurements were performed on the T2 generation of plants with 15-25 plants for each independent line. For flowering phenotype analysis, primary transformants were selected for the transgene and sown on soil and transferred to LD conditions after stratification (4°C, 3 days). Flowering time was determined in randomly distributed plants according to number of rosette leaves at the time of bolting (10 plants for wild type, *lux-4*, *lux-4* p*LUX*::*LUX*^*R146A*^ and *lux-4 pLUX::LUX*).

### Plasmid construction and generation of transgenic plants

For the *lux-4* p*LUX*::*LUX*^*R146A*^, *lux-4 pLUX::LUX* constructs, a ~800bp upstream fragment of *LUX* was PCR-amplified from genomic DNA. Full length CDS of *LUX* and *LUX*^*R146A*^ were PCR-amplified from the pESPRIT002 expression vector containing the respective CDS with an N-terminal FLAG tag added (SI Figure 2). NEBuilder^®^ HiFi DNA Assembly Kit (E2621S, NEB) was used for assembling the promoter fragment with the appropriate cDNA fragment, FLAG tag and vector backbone (pFP101 containing the *At2S3* promoter driven GFP for selection of transformants)(57). For a list of primers see Supplemental Table 1. Transgenic plants were generated by *Agrobacterium*-mediated gene transfer using the floral dip method (58). *Lux-4* plants were dipped with *Agrobacterium* containing *pLUX::LUX*^*R146A*^ or *pLUX::LUX* constructs to obtain *lux-4 pLUX::LUX*^*R146A*^ and *lux-4 pLUX::LUX* plants.

### RNA isolation and quantitative PCR

Plants were grown under short day conditions (8L:16D) for 7 days in 0.5 MS media and samples were harvested in intervals of 4 hours. 8-10 seedlings were harvested for each line at each time point. Total RNA was extracted using RNeasy Plant mini kit (Qiagen) according to manufacturer’s instructions. Total RNA (1µg) was treated with DNaseI (Roche) qRT-PCR was done using iTaq^®^ Universal SYBR^®^ Green One-Step Kit from Bio-Rad following manufacturer’s protocol. For the list of primers see SI Table 1. Expression of *PIF4* in different plant lines was determined through qRT-PCR with *PP2A* used as a control. qRT-PCR measurements were performed with a Bio-Rad CFX connect Real-Time system. Quantification was performed with the relative –ΔCt method, using *PP2A* for normalization. All quantification and statistical analysis were performed using CFX Maestro^TM^ software (Bio-Rad).

